# Precise Lineage Tracking Using Molecular Barcodes Demonstrates Fitness Trade-offs for Ivermectin Resistance in Nematodes

**DOI:** 10.1101/2024.11.08.622685

**Authors:** Zachary C. Stevenson, Eleanor Laufer, Annette O. Estevez, Kristin Robinson, Patrick C. Phillips

## Abstract

A fundamental tenet of evolutionary genetics is that the direction and strength of selection on individual loci varies with the environment. Barcoded evolutionary lineage tracking is a powerful approach for high-throughput measurement of selection within experimental evolution that to date has largely been restricted to studies within microbial systems, largely because the random integration of barcodes within animals is limited by physical and molecular protection of the germline. Here, we use the recently developed TARDIS barcoding system in *Caenorhabditis elegans* (Stevenson et al., 2023) to implement the first randomly inserted genomic-barcode experimental evolution animal model and use this system to precisely measure the influence of the concentration of the anthelmintic compound ivermectin on the strength of selection on an ivermectin resistance cassette. The combination of the trio of knockouts in neuronally expressed GluCl channels, *avr-14*, *avr-15*, and *glc-1*, has been previously demonstrated to provide resistance to ivermectin at high concentrations. Varying the concentration of ivermectin in liquid culture allows the strength of selection on these genes to be precisely controlled within populations of millions of individuals, yielding the largest animal experimental evolution study to date. The frequency of each barcode was determined at multiple time points via sequencing at deep coverage and then used to estimate the fitness of the individual lineages in the population. The mutations display a high cost to resistance at low concentrations, rapidly losing out to wildtype genotypes, but the balance tips in their favor when the ivermectin concentration exceeds 2nM. This trade-off in resistance is likely generated by a hindered rate of development in resistant individuals. Our results demonstrate that *C. elegans* can be used to generate high precision estimates of fitness using a high-throughput barcoding approach to yield novel insights into evolutionarily and economically important traits.

## Introduction

The interplay between environmental context and the direction and strength of selection on individual loci has been the fundamental underpinning of evolutionary genetics since its inception. Most standard population genetic models are grounded in two essential features: (1) that selection is relatively constant over time and (2) that selection can be best estimated in an aggregate fashion across all alleles with similar phenotypic outcomes. However, populations are never static—new mutations constantly arise and the environment is ever changing—meaning that the pattern of selection on individual loci is likely to change often, sometimes dramatically so (Bell, 2010). If there is a temporal or spatial structure to these environmental differences, then this variation can potentially lead to the maintenance of genetic diversity (Abdul-Rahman et al., 2021). Several examples of this process exist for antibiotics, small molecules, and more complex traits such as stress tolerance (Rudman et al., 2022). While initial formulations of population genetics focused on allelic change in a cross-sectional, generation-by-generation fashion, over the last several decades developments in molecular population genetics have shifted the focus on the importance of coalescence of evolutionary lineages for full inference of the evolutionary process, usually with a retrospective view (Wakely, 2016). The development of random barcoding approaches in microbial systems such as bacteria and yeast have allowed lineage-based approaches to be expanded into a fully experimental framework using a prospective approach (Ba et al., 2019; Blundell and Levy, 2014; Jahn et al., 2018; Jasinska et al., 2020; Levy et al., 2015). Yet, similar technologies have heretofore been missing for animals, largely because it is very difficult to transduce large libraries of DNA barcodes directly into animal gametes (Stevenson et al., 2023). We have recently developed a library-based transgenesis system called Transgenetic Arrays Resulting in Diverse Integrated Sequences (TARDIS) within the nematode *Caenorhabditis elegans* that overcomes this barrier in two steps: first by creating a diverse bar code library within the individual using an extra chromosomal array and then secondarily randomly incorporating individual bar code elements into a defined landing pad location via CRISPR/Cas9 activation in a subsequent generation (Stevenson et al., 2023). Here, we provide an exemplar for the application of TARDIS barcoding within experimental systems by exploring potential trade-offs in natural selection for resistance across a gradient of concentrations of the anthelmintic ivermectin, demonstrating that lineage-based approaches in experimental evolution can serve as a powerful means of generating high-precision estimates of the magnitude of natural selection in the face of environmental variation.

Insecticides represent a wide class of compounds that disrupt essential biological functions of insect pest populations and are widely used to improve health outcomes for humans and animals, as well as to support agricultural systems (Araújo et al., 2023). In natural populations resistance to pesticides has routinely evolved (Bras et al., 2022; Hawkins et al., 2019; Shi et al., 2019; UK et al., 2014), and the acquisition and spread of insecticide resistance has long served as an important exemplar of evolution within natural populations (Crow, 1974; ffrench-Constant, 2013; Freeman et al., 2021; Mallet, 1989; Pu and Chung, 2024). Indeed, the evolution of insecticide resistance is a major concern due to its potential global economic impact (Forgash, 1984; Mallet, 1989; Pimentel, 2005; Robinson, 2002). For example, in the United States, it has been estimated that resistance to insecticides costs over $10 billion annually (Gould et al., 2018).

Ivermectin, the most widely used anthelmintic drug (Campbell, 1993; Geurden et al., 2015; Gill et al., 1991; Leathwick et al., 2012; Prichard, 2007; Shoop, 1993) is used worldwide for controlling nematode infestations both within livestock and companion animals and in humans for the treatment of crippling parasite diseases such as ascariasis, which can infect the lung and intestines, and onchocerciasis, which causes river blindness (Conterno et al., 2020; Leung et al., 2020; Sulik et al., 2023). Rapid development of resistance to ivermectin in particular is a growing problem (Doyle et al., 2022). Ivermectin works by activating the glutamate-gated chloride (GluCl) channels, leading to hyperpolarization (Ardelli et al., 2009; Dent et al., 1997). Within laboratory populations of *C. elegans*, these neurological effects can be quantified by measuring the rate of muscle-based phenotypes, such as pharyngeal pumping (Weeks et al., 2018). Extensive screening efforts have identified three mutations in the loci encoding GluCl channels (*avr-14*, *avr-15*, and *glc-1*) that, in combination, lead to a roughly 4000X increase in resistance to ivermectin (Dent et al., 2000; Shaver et al., 2024). The combination of the underlying functional biology of these mutants and the power of *C. elegans* as a system for experimental evolution (Teotónio et al., 2017) makes this an especially powerful approach to address the question of adaptive mutations for ivermectin resistance in nematodes.

Here, we report the first-ever randomly barcoded evolutionary lineage tracking experiment performed within an animal system, *Caenorhabditis elegans*, which allows replicated measurements of selection coefficients of a known mutant within a well-defined environmental context. We barcode populations of *C. elegans* utilizing TARDIS—a high-throughput transgenic methodology (Stevenson et al., 2023)—with unique collections of barcodes to distinguish between wildtype and mutant backgrounds (Figure 1A). We also present a modified liquid culture protocol for growing several multi-million sized animal populations in parallel (Figure 1B, Figure 1–supplemental figure 1), making this experiment, to our knowledge, the largest animal experimental evolution study conducted to date. Utilizing barcode sequencing upon each transfer (Figure 1C), our results show that selection is dependent on the concentration of ivermectin in the liquid environment, illustrating a clear fitness trade-off for the mutants depending on environmental conditions and which manifests phenotypically as a trade-off in developmental rate. Our project serves as an initial exemplar of lineage tracking within an animal context, which can be applied generally to study lineage dynamics in experimental populations.

**Figure 1.**
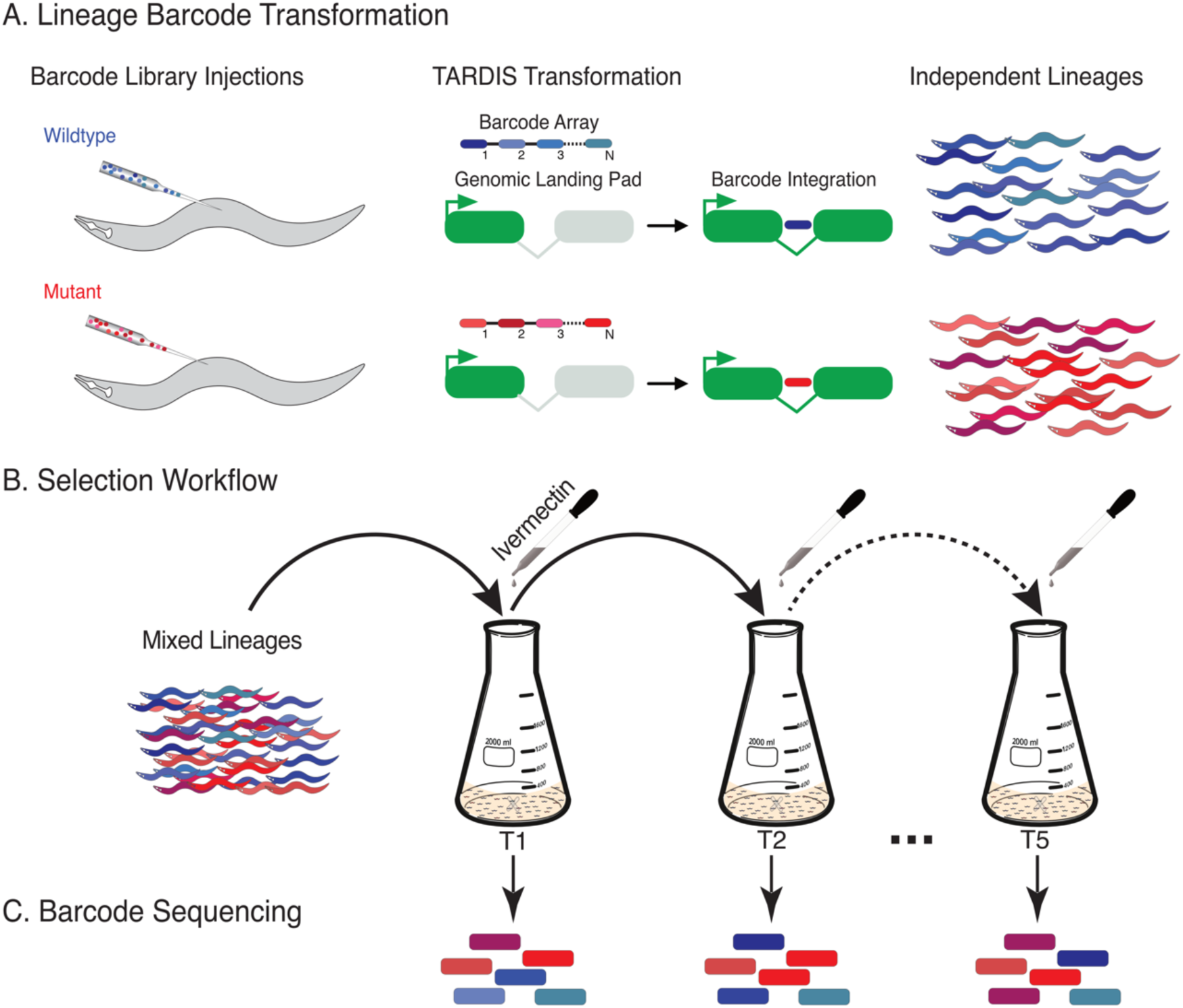
Experimental overview. A) Lineage transformation following TARDIS transgenesis. Barcodes are integrated within synthetic introns for hygromycin B resistance and were engineered to contain ‘constant’ bases to distinguish from which genetic background lineages originated. B) 30-60 lineages were then pooled and serially cultured in various concentrations of ivermectin (0nM to 5nM with 1nM increments) for a total of five transfers (T1-T5). C) At each transfer, a portion of the population was lysed and genomic DNA was extracted. Barcodes were amplified and quantified by NGS.

**Figure 1 – figure supplement 1.**
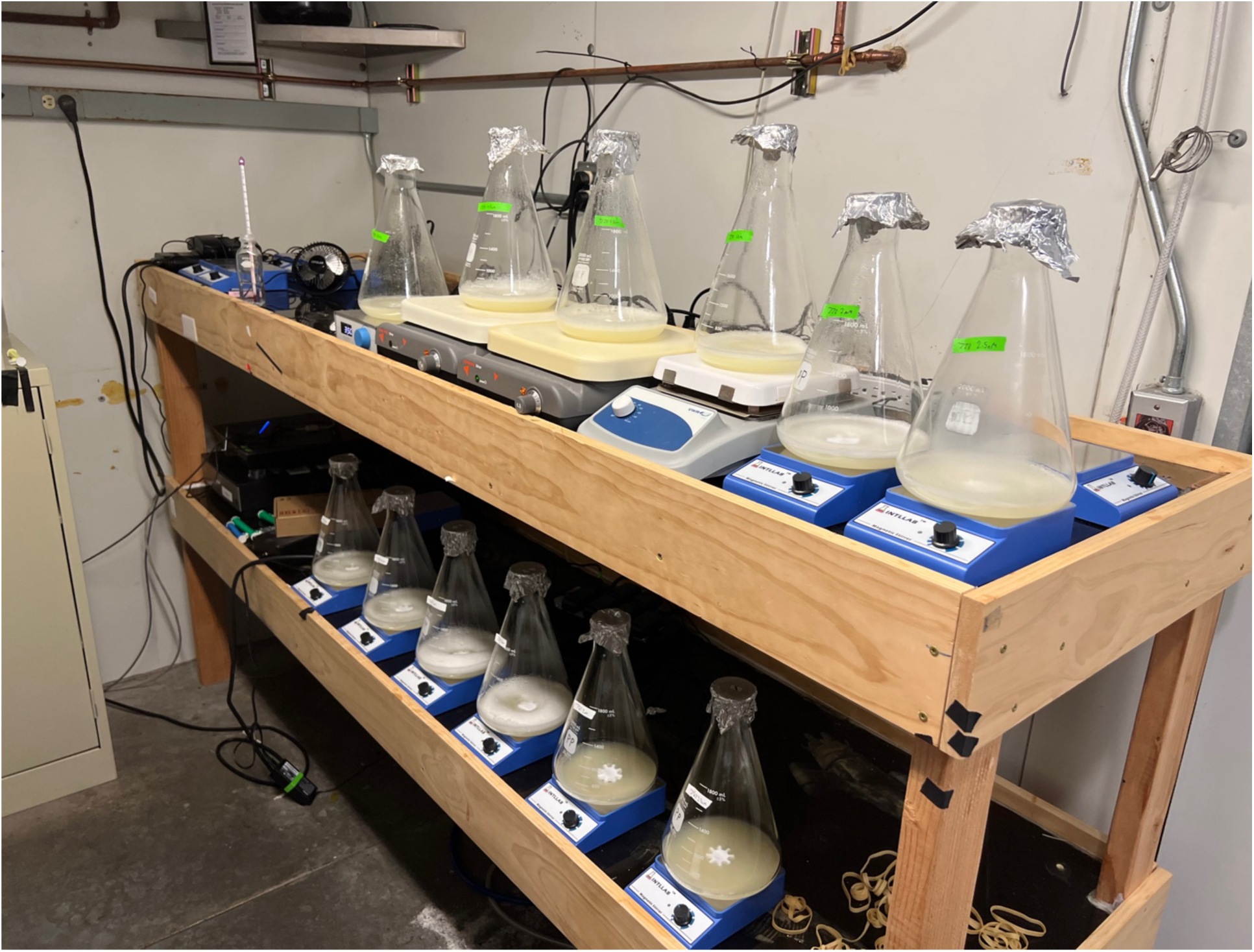
Photograph of the liquid culture environment in our temperature-controlled unit.

## Results

### Ivermectin exposure creates environmentally dependent and dynamic selection across generations

We performed a competition experiment with multiple barcoded lineages of both wildtype and mutant backgrounds in large scale liquid culture and several concentrations of ivermectin exposure. In this way, the mutations and wildtype individuals are “identical by kind” as determined by their allelic state, but only individuals within a given barcoded lineage are “identical by descent” (Lewontin, 1986). Cultures were serially transferred a total of five times (T1-T5) with barcode frequencies measured at each transfer (Figure 2), allowing us to access the density of each lineage within the populations. Census size populations were generally maintained above 10^5^ and often surpassed 10^6^ individuals (Figure 2–supplementary figure 1), greatly beyond the threshold for drift to have influenced our results. Overall, we observed a gradient of selection overtime, with lower [ivermectin] favoring the wildtype and higher concentrations favoring the triple mutant (Figure 2).

**Figure 2.**
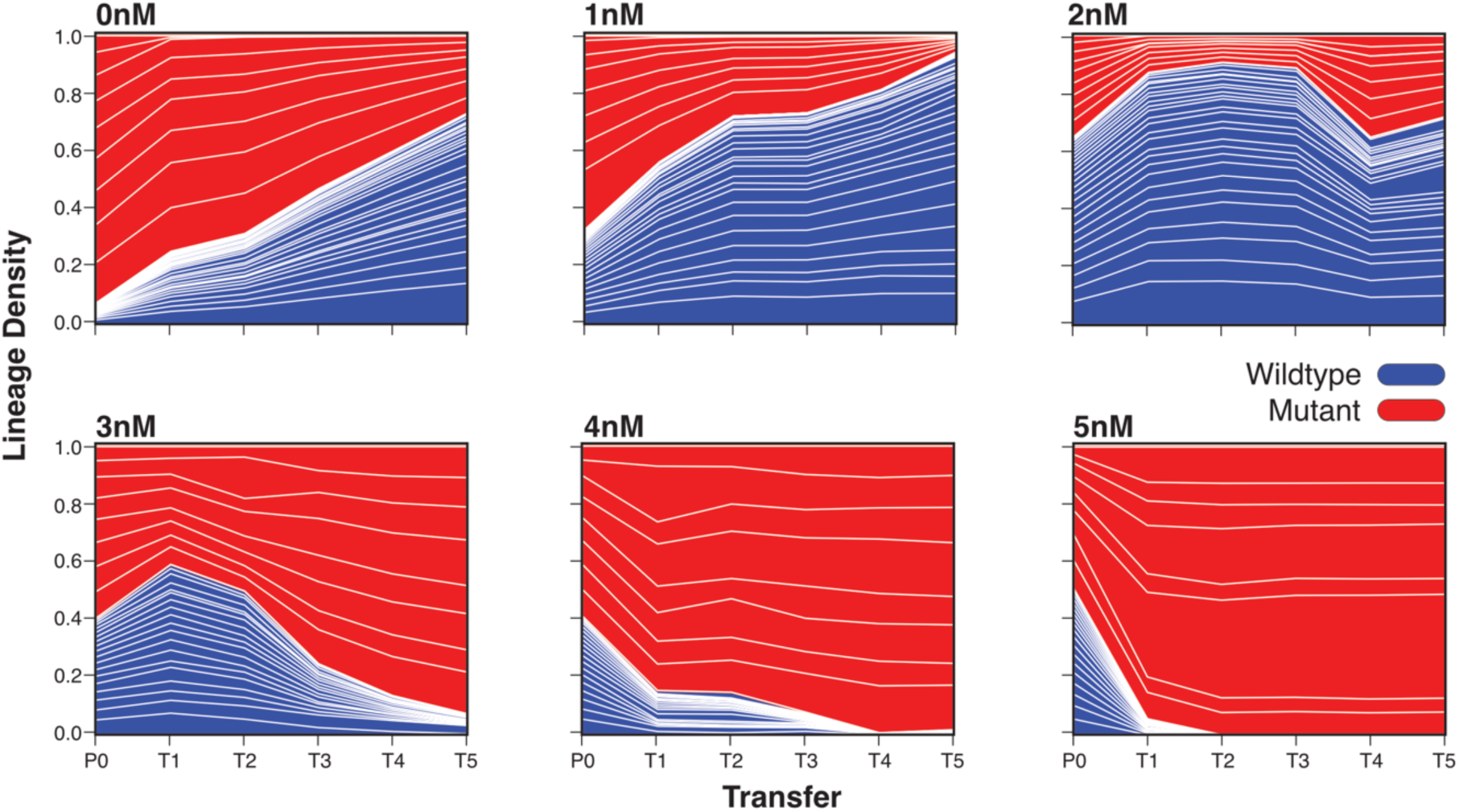
Example lineage frequencies across ivermectin conditions. Transfers (T1-T5) are denoted along the x-axis with the parental generation (P0). Lineage density as measured by the barcode frequency is denoted on the y-axis. At P0, wildtype (blue) and triple mutant (red) lineages were combined at different starting frequencies that varied with [ivermectin]. For 0nM and 1nM we see a clear trend towards a wildtype advantage–there is a cost for being resistant to ivermectin. In the 2nM condition we start to see the wildtype lineage receiving less of an advantage. For 3nM, 4nM, and 5nM we see a trend towards increasing mutant frequency for each condition.

**Figure 2 – figure supplement 1.**
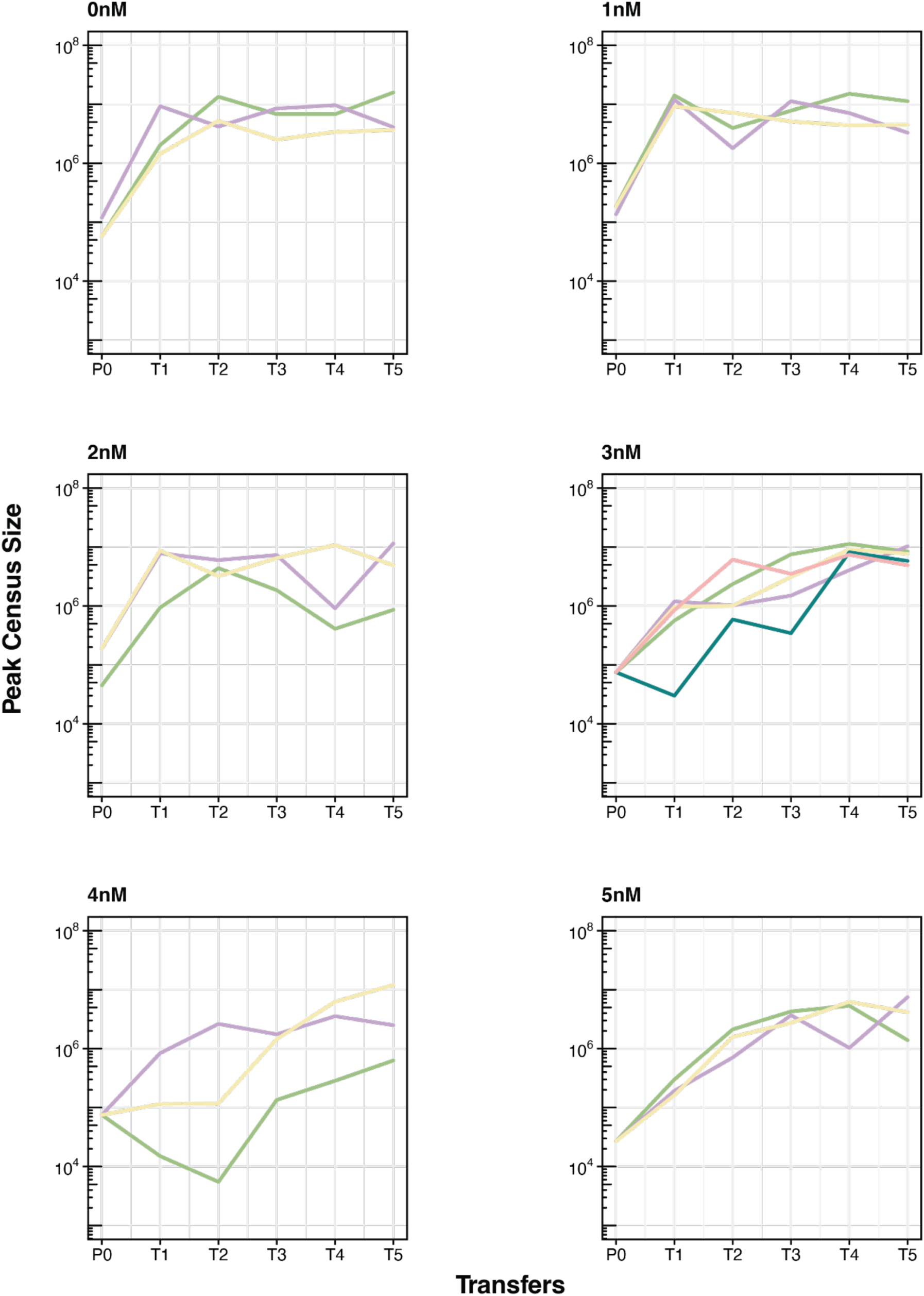
Peak census size for each replicate population plotted at log_10_. Colors indicate unique replicates within a concentration. Peak population sizes often surpass a million individuals.

In our control (Figure 2, 0nM), the wildtype background was favored suggesting that without selection pressure there was a significant deleterious cost associated with the triple mutant background. At the lowest concentrations of ivermectin (Figure 2, 1nM), the advantage of the wildtype background remains, however, it is lessened in comparison to the 0nM condition. With increasing concentrations of ivermectin, a transition around the 2nM mark occurs whereby the mutant becomes increasingly favored with each stepwise increase in [ivermectin] (Figure 2, 3nM to 5nM). Interestingly, selection was not constant and, in some cases, shifted from transfer to transfer (see Figure 2, 1nM T2 to T3 and 2nM T4 to T5). This variability, combined with the early wildtype advantage in some cases (Figure 2, 2nM and 3nM, P0 to T1), clearly suggests that multiple generations should be used to accurately measure fitness when performing competition experiments. Among our lineages, we clearly saw each lineage following similar trajectories to the other lineages within the same genetic background. This provided an internal quality control since there were no adaptive mutations of large effect occurring in the background which would generally be invisible in standard competition experiments.

### Mutant selection coefficients increase with ivermectin concentration

Utilizing the barcode frequencies, we were able to estimate the selection coefficients of each lineage from the mixed populations after the final transfer (T5). Each lineage acted as a separate measurement of the strength of selection and provided internal replication and measured variation in the selection coefficient for that population. With the wildtype selection coefficients held constant at zero, selection on the triple mutant rose exponentially alongside [ivermectin] with a strong correlation (adjusted *R*^2^ = 0.94, *F*2,149, *p* < 2.2×10^-16^) (Figure 3). Without ivermectin (0nM), the triple mutant was conditionally deleterious with a selection coefficient of approximately *s* ≈ -0.5. Thus, there was a clear trade-off conferred in the absence of ivermectin. By simply increasing the ivermectin concentration, we were able to lessen the selective pressure and swap the deleterious and advantaged backgrounds. Similar to our lineage trajectories, the wildtype genotype was favored under 2nM, with a point of neutrality at approximately 1.5nM. Generally, we observed large shifts in the selection coefficient per condition. Additional transfers were essential for accurately estimating the selection coefficients, as we generally saw fluctuations in the transfer early on (Figure 2). With additional, intermediate, concentrations of ivermectin, and larger number of transfers, even finer resolution of selection coefficients could possibly be achieved.

**Figure 3.**
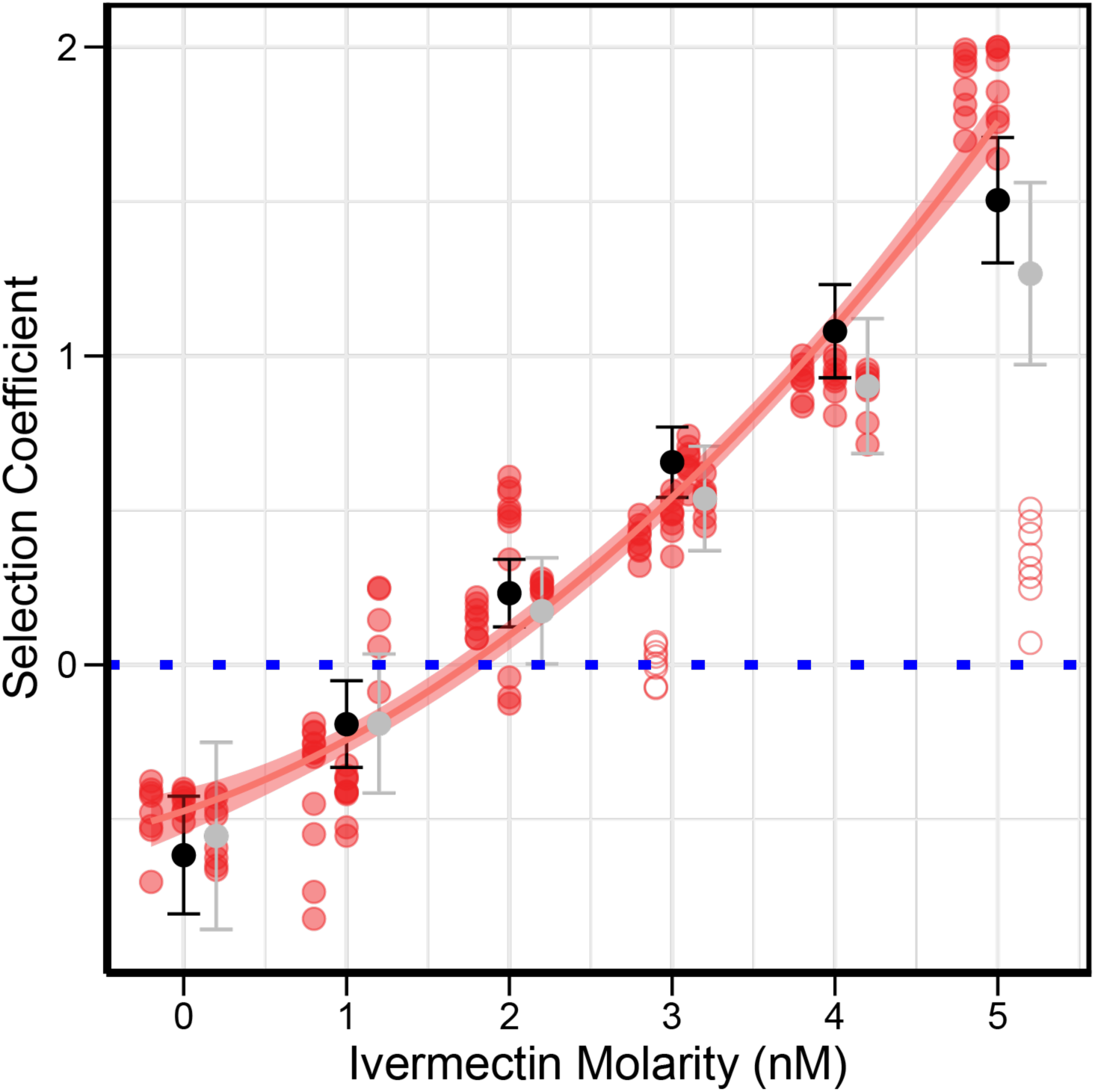
Mutant selection coefficient in relation to ivermectin concentration. Blue dotted line represents the wildtype selection coefficient, which was normalized to zero for each concentration. Each selection condition has three replicates, 3nM has an additional two replicates. Individual replicates are jittered into columns within an ivermectin concentration. Hollow circles represent outlier replicates circles that were excluded from curve fitting. Black bars represent the least mean squared confidence intervals for each concentration. Gray bars include the outlier replicates. Fitted polynomial is S=0.053[ivermectin]^2^ + 0.18[ivermectin] - 0.5, adjusted R^2^ = 0.94, p-value=<2.2×10^-16^. We see a clear exponential increase in fitness with concentration for the mutant lineages.

### Ivermectin resistance generates a trade-off on developmental rate

While maintaining JD608 (triple GluCl mutant) and N2 (WT) strains, we observed a noticeable delay in development for the mutant strain compared to wildtype in the absence of ivermectin. In contrast, when grown on plates in the presence of ivermectin, we observed that wildtype worms were significantly delayed, leading us to hypothesize that delayed development, and a concomitant delay in reproduction, could be the source of selective tradeoff to ivermectin. To test this, we hypochlorite synchronized a single barcoded lineage from both the wildtype and mutant backgrounds and exposed them to ivermectin in a liquid culture environment that paralleled the conditions that we used for the selection experiment. At two separate time points (72 and 96 hours post synchronization), we took samples from the culture and counted the total number of worms, and what proportion had reached adulthood (Figure 4, 72 hours post synchronization, Figure 4–figure supplement 1, 96 hours post synchronization). We found there to be a strong correlation with reduced developmental rate and ivermectin concentration (72 hours, *p* < 2.2×10^-16^). At 0nM, the development of the animals with the wildtype background outpaced the development of those with the mutant background. This advanced development likely provides a significant adaptive advantage to the animals with the wildtype background when compared to the triple mutant animals competing in the same environment. However, when animals are grown in increasing ivermectin concentrations, wildtype development is “stunted.”

**Figure 4.**
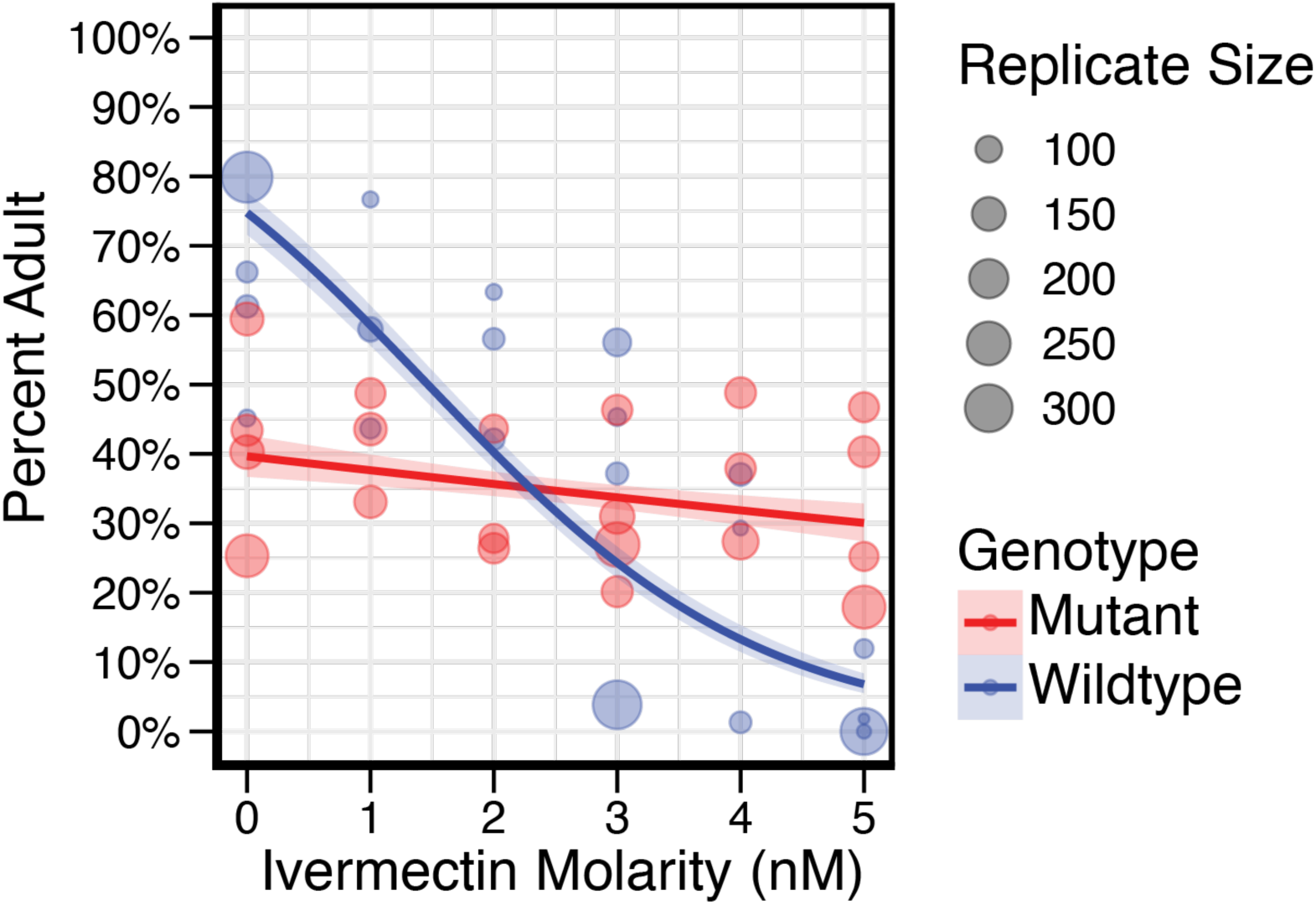
Developmental delay at 72 hours while developing in ivermectin. Ratios represent the number of adults over the total number counted. Wildtype is represented in blue, while mutant is represented in red. Total worms counted per replicate is represented by circle size. We see a clear impact of ivermectin on development (p- value=<2.2×10^-16^). Shading denotes 95% confidence interval. We see the wildtype background is more advanced in development compared to the mutant background at the 0nM concentration up until approximately 3nM, where the mutant shows a developmental advantage. At all five conditions we see the mutant background is developing at relatively the same rate.

**Figure 4 – figure supplement 1.**
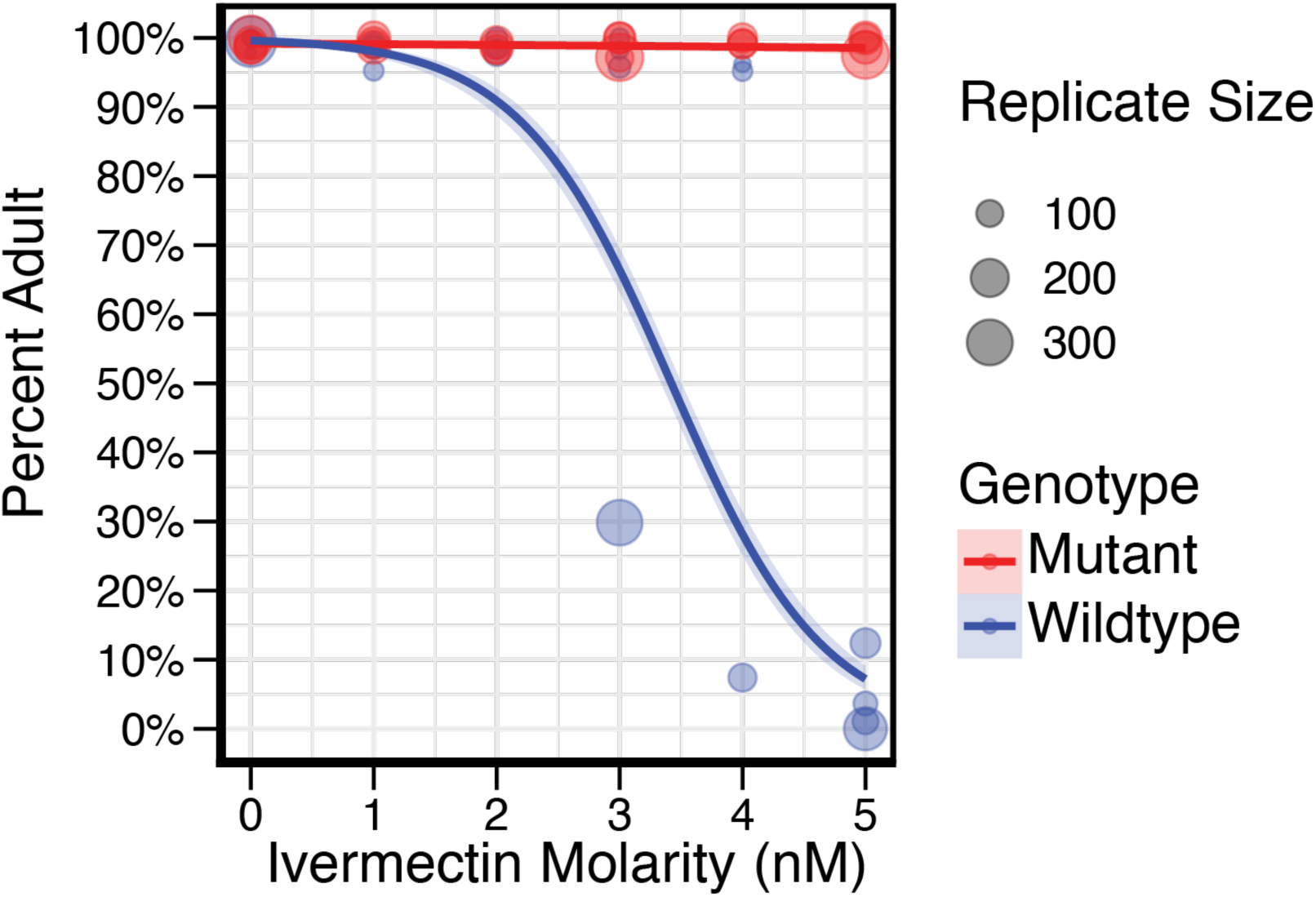
Developmental delay at 96 hours while developing in ivermectin. Wildtype continues to develop slowly in deleterious concentrations of ivermectin, however, it is progressing in 3nM and 4nM. At 5nM, the wildtype background remains highly stunted with very few individuals reaching adulthood.

Development of the wildtype background animals when grown in 3nM ivermectin appears to be distributed across several larval stages, with adults making up a significantly smaller proportion of the total progeny numbers when compared to those grown without ivermectin (Figure 4–figure supplement 2). Additionally, when compared to the mutant background animals grown in the presence of 3nM ivermectin, there was no significant difference in the percentage of adults between the two strains. At our highest concentration of ivermectin, 5nM, we observed very few adults from the wildtype background animals (3.4% compared to the 63.1% observed at 0nM, 72 hours). In contrast, the mutant background animals developed at approximately the same rate across all ivermectin concentrations although there was a slight initial delay at 72 hours that was not observed after 96 hours (Figure 4 compared to Figure 4–supplement 1). Overall, then, the developmental delay hypothesis is strongly supported, with a clear trend towards slower development for the wildtype strain on increasing concentrations of ivermectin. Remarkably, we observe a similar crossover point for wildtype vs. mutant success of approximately 2.5nM for both the developmental delay and selection estimates.

**Figure 4 – figure supplement 2.**
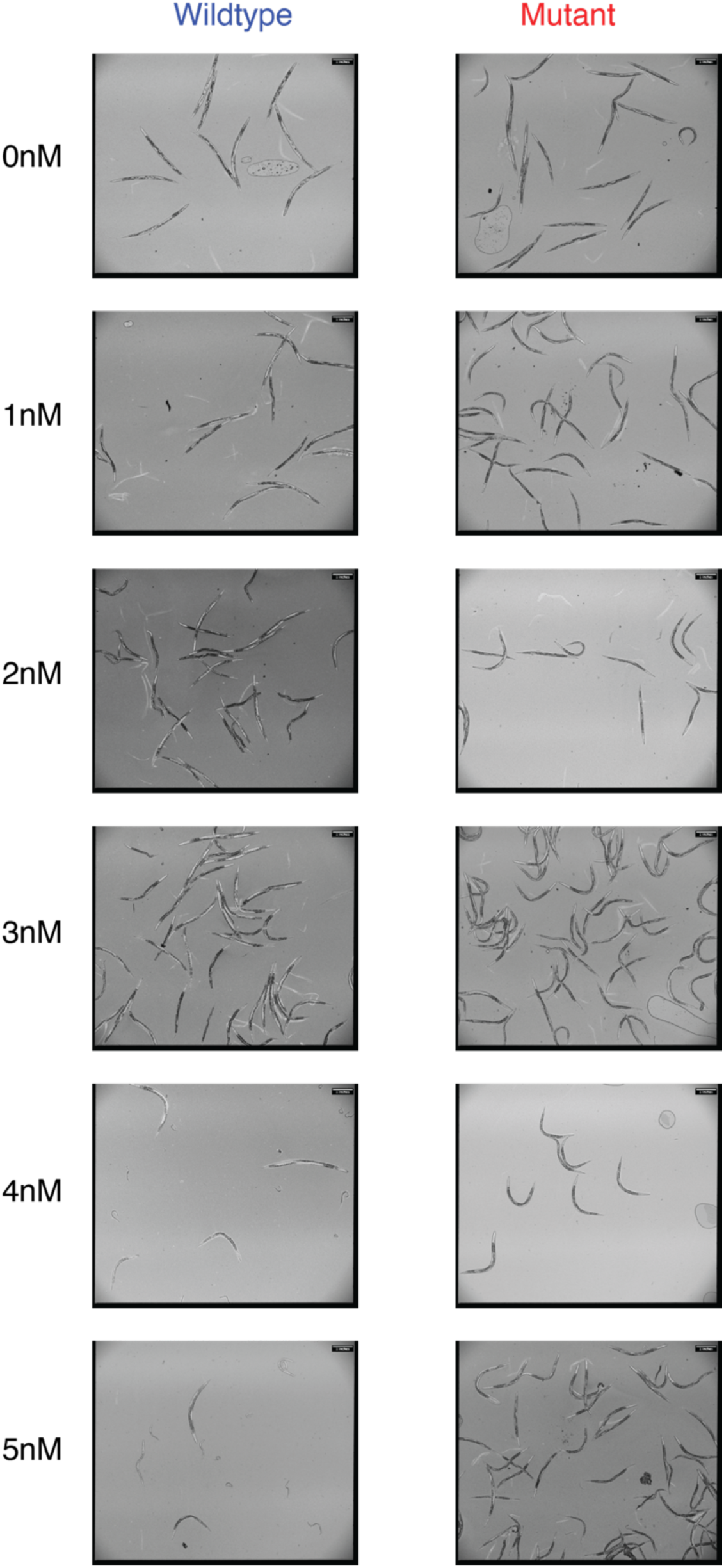
Representative images of populations at 72 hours while developing in ivermectin. For wildtype, below 3nM, development is approximately similar with noticeable heterogeneity in developmental stage at 3nM. After 4nM, there is a steep decline in development. The mutant strain remains consistent in development across the ivermectin concentrations.

## Discussion

Darwin (1859) is most recognized for introducing the idea of natural selection, but it is really his overall vision of “descent with modification” that captures the entire scope of the evolutionary process. Full reconciliation of phylogenetic, molecular evolutionary, and population genetic perspectives on evolutionary relationships depends on being able to tie microevolutionary processes to lineage-based estimates of evolutionary change. Here we present the first barcoded evolutionary lineage tracking experiment performed within an animal system. Our results demonstrate that we can precisely and reproducibly measure selection within a specific environmental context and change the evolutionary advantage of a given haplotype by changing the environment in which it is found. Utilizing our liquid culture approach with the nematicide ivermectin as the selective agent, we were able to grow populations in the many millions, creating the largest experimental evolution study conducted within an animal system to date. The application of our new unique random barcoding system used here both provides interesting insights into an important agricultural intervention and paves the way for the application of this technology to a wide set of important evolutionary questions.

### Naturally occurring genetic variation and hypothesis testing of ivermectin resistance

Resistance to ivermectin has already been widely observed within natural populations of nematodes (Hawkins et al., 2019). Likely, as the application of ivermectin cannot be maintained at a constant level, parasites are exposed to below-therapeutic concentrations of ivermectin in either missed doses or incomplete treatments, providing an opportunity for ivermectin-resistant mutations to increase in frequency in the population before they can be eliminated via a lethal dose (Fissiha and Kinde, 2021). We used a system of three synthetic resistance mutations to establish the framework for experimental evolution and barcode lineage tracking used here. While this system is certainly artificial to a degree, in the future “parasitized” strains of *C. elegans* in which the native allele is swapped for a resistance allele from a natural parasite could provide further insight into the evolution of anthelmintic resistance utilizing a non-parasitic lab model (Zamanian and Andersen, 2016). Within natural nematode populations, potential resistance alleles in *avr-14* and *avr-15* have yet to be found (Doyle et al., 2022), however there is evidence for selective mutations within *glc-1* (Ghosh et al., 2012), as well as in other *C. elegans* homologs such as the *cky-1* mutation found in the parasitic nematode *Haemonchus contortus* (Doyle et al., 2022). There is also evidence of other potential ivermectin resistance genes within *C. elegans* whose effects could be quantified more precisely using the methods developed here (Hunt et al., 1994; Su and Dent, 2015).

We used all three mutations in tandem for this study, but because we were able to measure selection in the triple mutant background, it should be possible to barcode individual strains with single or double mutations in *avr-14*, *avr-15*, and *glc-1* to discover possible adaptive intermediates across a range of concentrations. Shaver et al. (2024) have recently competed individual strains with new knockout alleles for *avr-14*, *avr-15*, and *glc-1*, which are not the conical alleles used in our study. In their study, the selection and control conditions show results that differ somewhat from our study. For the single selective condition they used (1.5nM), the the triple mutant jumped from a 10% allele frequency to approximately 100% within two generations, suggesting that selection was extremely strong under their conditions. While we did not specifically investigate 1.5nM, from our dosage curve we would predict 1.5nM to be approximately neutral, with possible bias towards wildtype. Shaver et al. (2024) also observed a much stronger deleterious condition in their control, DMSO, where after a single generation the triple mutant plummeted from approximately 50% to below 10%. The differences in these outcomes suggest that particulars in experimental conditions are likely to matter. For example, Shavers et al. (2024) defined a generation as approximately seven days, where as our transfers are every five days. If we compare the end point of our experiment at 25 days to their 3^rd^ generation (approximately 21 days), we might be able to explain a portion of the drastic selective difference in the control condition (DMSO). Perhaps most importantly, Shavers et al. (2024) performed their selection experiments on plates whereas we used large scale liquid culture, and they investigated a single concentration whereas we estimated the entire response function for ivermectin resistance. The specific contribution of functional interactions among these genes to the pattern of natural selection we observe here certainly merits further investigation.

### Building upon the barcoded lineage tracking approach

While this project implements the analysis of randomly barcoded lineages within an animal system for the first time, microbial systems using similar approaches have been well developed for evolutionary studies, particularly for estimating the distribution of fitness effects (Ba et al., 2019; Blundell and Levy, 2014; Levy et al., 2015). For our system, adoption of several, highly diverse barcode TARDIS libraries could reasonably result in several hundred thousand unique lineages (Stevenson et al., 2023), comparable to even the largest barcoded experiments performed in microbes. While population sizes within the billions would be unrealistic, populations within the several hundred million are possible by simply scaling the liquid culture system developed here. Similar to microbial systems, we capitalized upon the self-fertilizing nature of hermaphroditic *C. elegans* to ‘lock’ a barcode within a lineage, mimicking asexual reproduction used in microbial lineage tracking experiments. However, barcoding under sexual reproduction could be feasible by barcoding across multiple haplotypes of a chromosome and could be applied to a variety of questions regarding sexual selection, sexual dimorphism, and adaptation (Kasimatis et al., 2021).

### Pleiotropic effects and trade-offs in adaptation

Pleiotropic effects of potentially beneficial mutations are thought to be widespread within genetic systems (Zhang, 2023). The consequences of pleiotropic effects can lead to adaptive trade-offs, where a mutation can provide a benefit within one environmental context and not within another (Bakerlee et al., 2021; Giannattasio et al., 2013; Jerison et al., 2020; Roff, 1992; Schmidlin et al., 2024; Wang et al., 2015). A similar example of an adaptive trade-off occurs between the garter snake, *Thamnophis sirtalis*, and its prey, the newt *Taricha granulosa,* which produces a neurotoxin, tetrodoxin (TTX) as an antipredator defense (Brodie III and Brodie Jr., 1999, 1991, 1990). While resistance to TTX has evolved in the garter snake it is also accompanied by an adaptive trade-off in which garter snakes with resistant-mutations in the TTX-binding site of Nav1.4, a voltage gated-sodium channel, have impaired movement (Carlo et al., 2024). A physiological tradeoff with resistance at the level of neuronal signaling of this kind appears to mirror the results seen here, in which we find a clear adaptive trade-off between ivermectin resistance and developmental rate in our experimental populations. A trade-off of this kind could potentially help drive the dynamics of natural resistance to ivermectin in the field, and indeed, when exposed to increasing concentrations of ivermectin the parasitic nematode *Haemonchus contortus* shows decreasing rates of larval development (Tuersong et al., 2022).

Experimental evolution in the laboratory, where selective and environmental conditions can be finely controlled, can be used to develop precise hypotheses that can be tested within natural populations where the complexity of mitigating factors might often confound our ability to cleanly addressing a specific functional hypothesis.

### Advantages and applications of multi-lineage barcoding in assessing mutant fitness and evolutionary dynamics

Strictly speaking, our random barcoding approach is not absolutely necessary for this work, as it is possible to assess mutant allele frequencies directly, or by using a simple co-marker such as fluorescence or molecular probes (Kasimatis et al., 2022; Murray and Cutter, 2011; Shaver et al., 2024; Webster et al., 2022). There are several distinct advantages to testing multi-lineage barcodes of the same allelic set, however. First, each lineage provides a replicated estimate of fitness in a given environment and trial. This allows a level of precision and statistical rigor that would otherwise be impossible. In our case, we found that lineage-specific fitness estimates tended to be very similar and to provide excellent estimates of mutant fitness. The interesting exception is when the concentration of ivermectin was right at the trade-off balance point (∼2nM). Here we saw a substantial increase in among-lineage variance within each replicate, as might be expected when a haplotype is near the neutral threshold. So, in this case, replicated lineage estimates are essential for providing high precision estimates of fitness, even when the selection coefficient is near zero. We also observed a few replicates in which the ivermectin addition and/or response were clear outliers (*e.g.,* a single replicate in both 3nM and 5nM both showed large deviations from the other replicates). Yet, since all the lineages responded in the same aberrant way—in a manner that was completely inconsistent with the entire experiment— we felt confident that the entire replicate had an unknown error during the execution of the experiment (possible causes could be accidental misapplication of ivermectin, a contaminate in the flask which persisted across the replicate, or microenvironmental changes which could impact the overall response to ivermectin) rather than being part of “normal” sampling variance across genotypes. However, it is important to note that our overall conclusions are completely unchanged if these replicates are included in the analysis, as only minor quantitative details of the response function are altered (Fig. 3). Second, having replicated lineages protects against a very serious confounding factor in experimental evolution studies: *de novo* background mutations. For example, if a new “high fitness” mutation arises spontaneously during the course of the experiment, it is impossible to separate its effect from the main effect of the mutant which is under study, especially when only assessing the allele frequency of the mutant itself. While we did not observe this in our current study, we have anecdotally observed this phenomenon while perfecting these methods, and it most certainly would be a caveat for experiments that run for longer durations than those presented here.

The ability to link novel mutations to unique lineages—and therefore unique evolutionary histories—is of course the real strength of barcoding based approaches. This has been extremely successful within a variety of single-cell systems, including bacteria (Jahn et al., 2018) and yeast (Ba et al., 2019; Blundell and Levy, 2014; Levy et al., 2015; Schmidlin et al., 2024), as well as in the proliferation of cancer cells (Lu et al., 2011). Our work establishes the groundwork for being able to conduct similar experiments in intact multicellular animals. The current study, while providing important insights into fitness trade-offs, also demonstrates that this barcoding can be used more generally. In establishing the TARDIS system, we showed that it is possible to generate several thousand barcodes via a single injection in a carrier that is a precursor to later lineage work. Combining a number of these precursors together, before barcode activation, allows the system to be scaled up into the hundreds of thousands needed for more general *de novo* mutation studies. So, in this way, this work illustrates how the next steps on that path can progress for studies centered on multicellular animals.

## Conclusions

In conclusion, we have presented the first barcoded lineage tracking animal experiment evolution, in what is also the largest animal experimental evolution study conducted to date. We created a simple experimental design to quantitatively measure selective contributions within a highly controlled environmental context and showed we can experimentally modulate the strength of selection, even changing the adaptive background, by changing the concentration of a simple small molecule drug ivermectin. We find there is an evolutionary cost to being resistant to ivermectin, which phenotypically manifests in delayed development in the absence of ivermectin. However, in the presence of ivermectin, we find sensitive individuals have highly stunted development, and therefore selection favors the resistant individuals. Our results thus highlight the kind of pleiotropic tradeoff that underlies many central ideas in evolutionary genetics, including the response of natural populations to human interventions such as insecticides and antibiotics.

## Materials and Methods

### Key reagents table

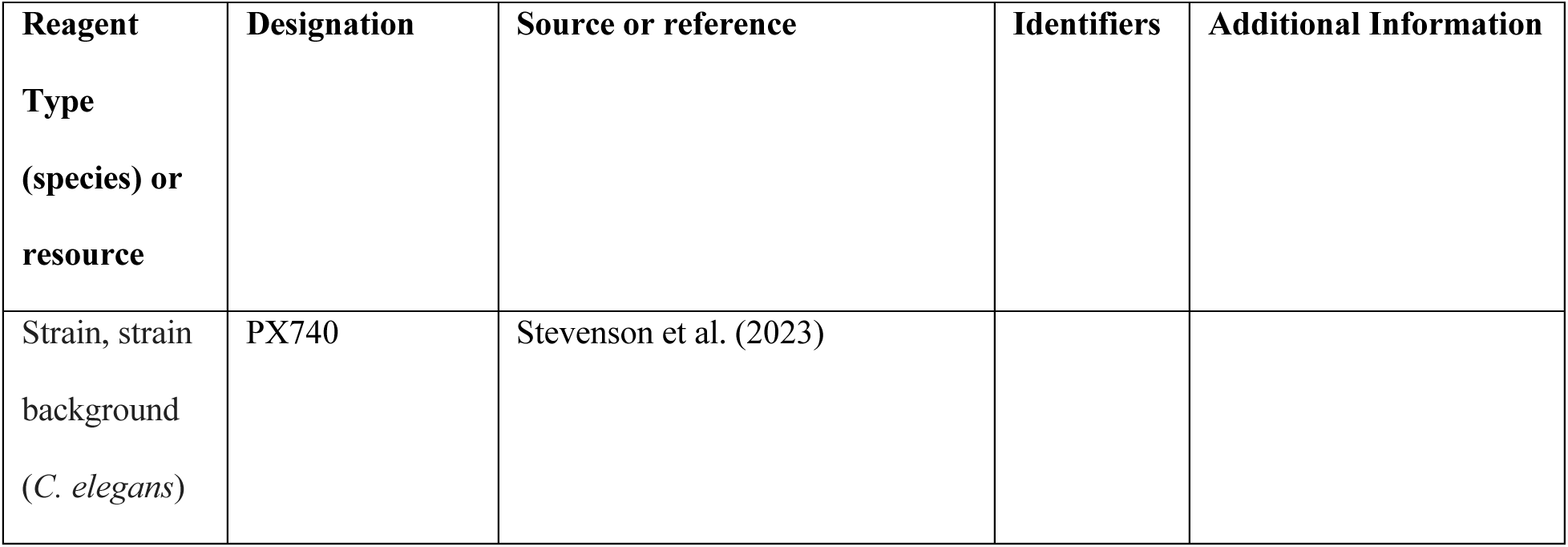
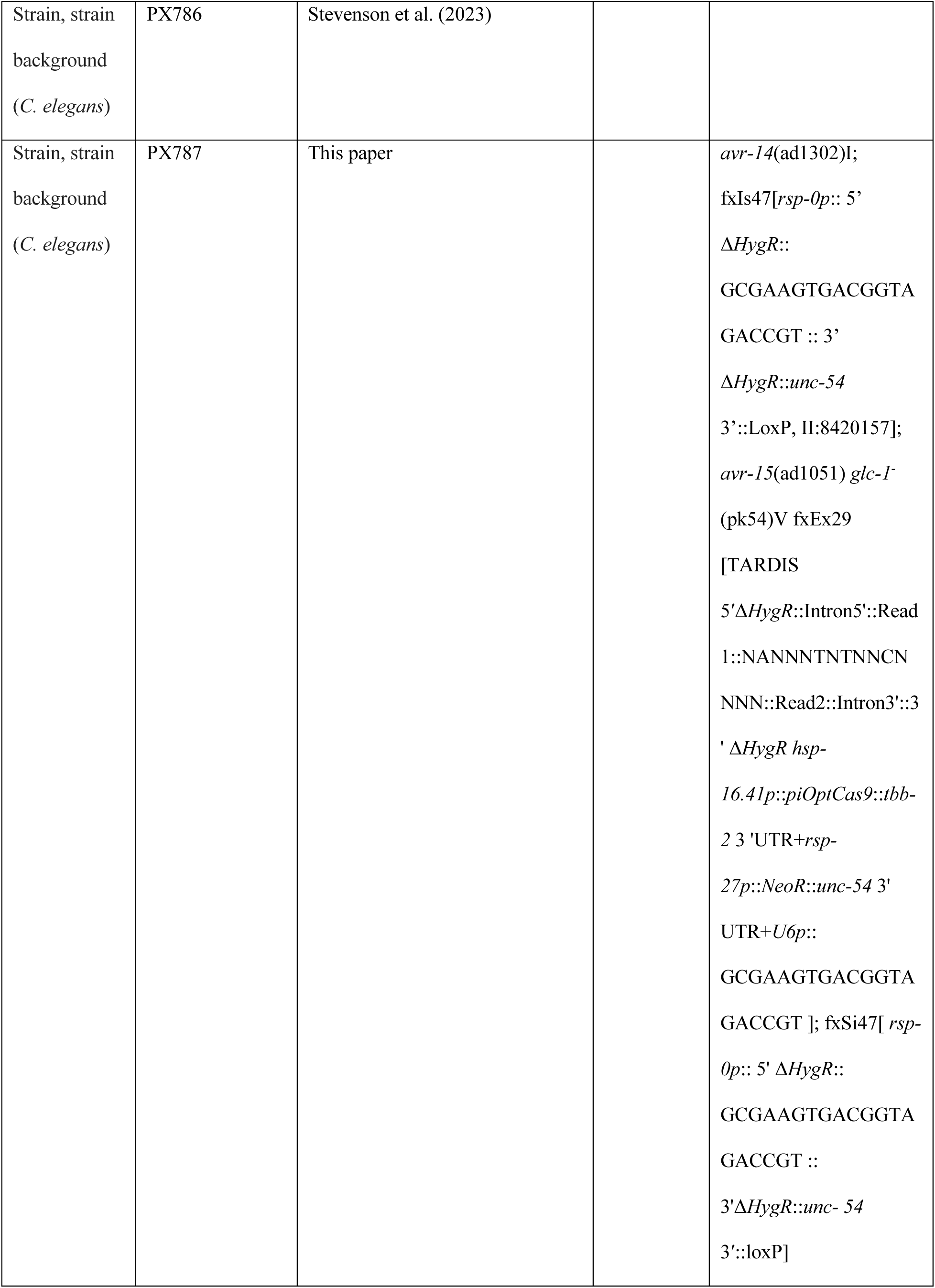
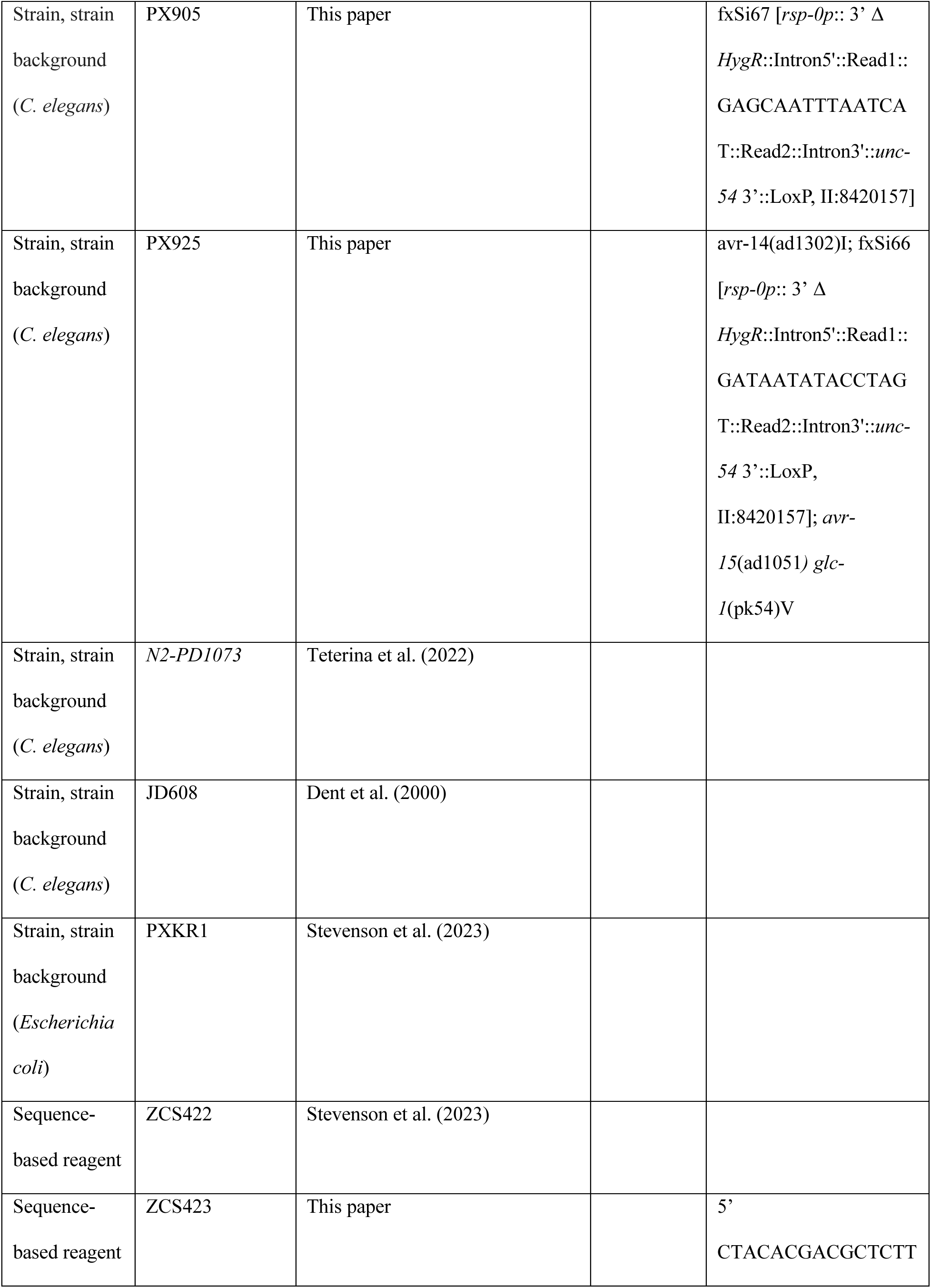
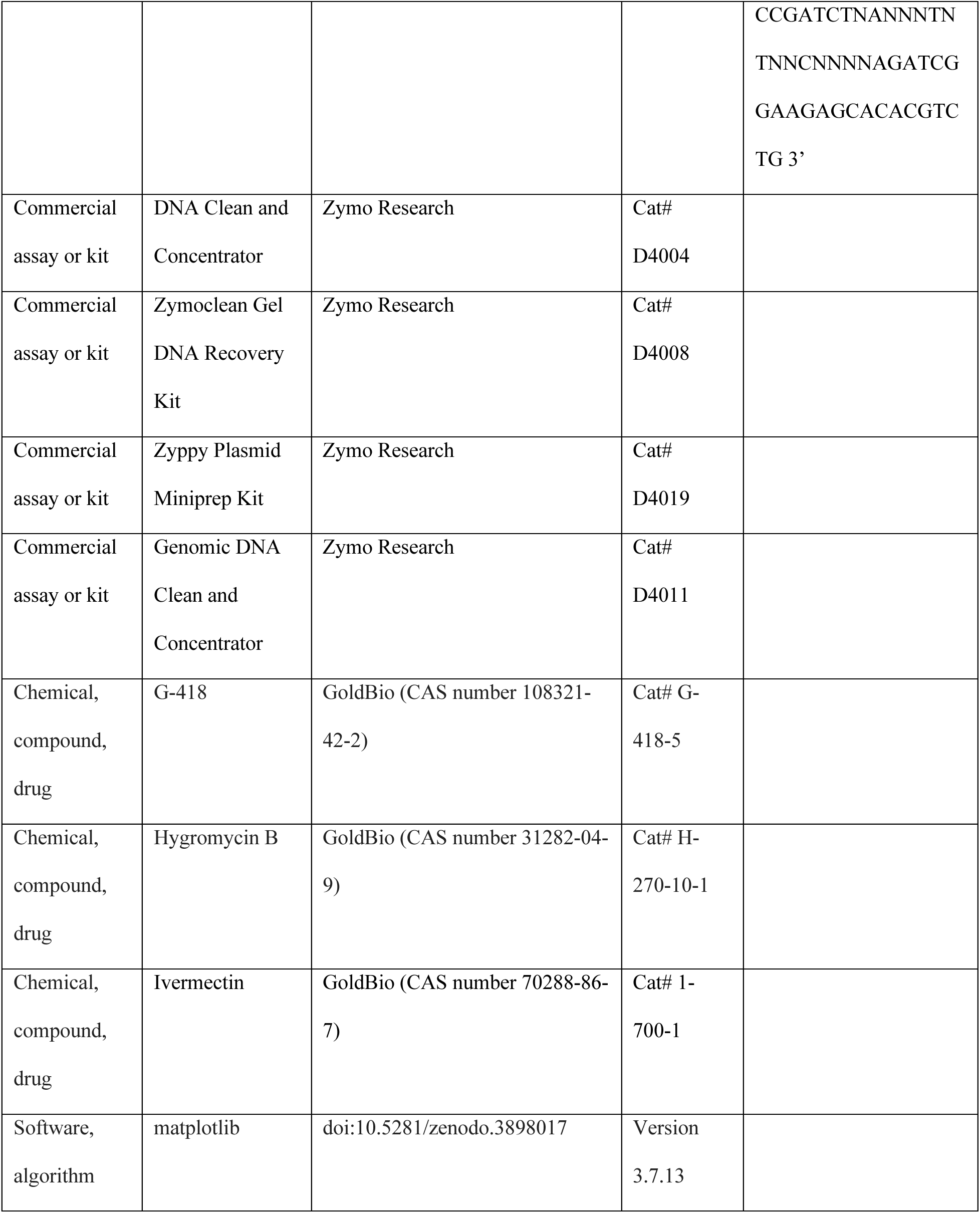
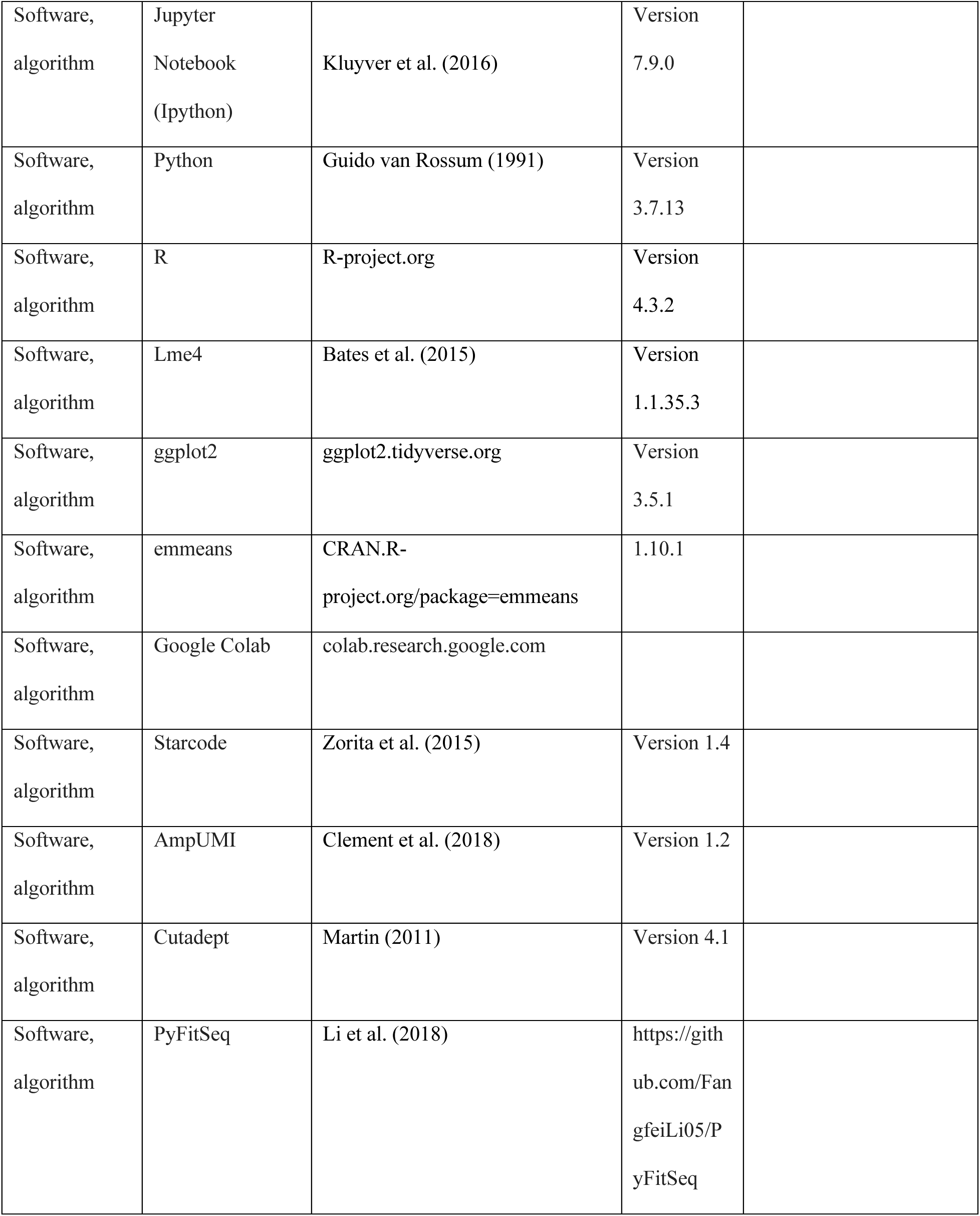

#### General reagents and C. elegans strain maintenance

Strain, plasmids, and reagents generated and utilized in this manuscript can be found in the key resource table. All plasmids have been described prior in Stevenson et. al. (2023).

Unless otherwise indicated, *C. elegans* strains were maintained at 20°C on nematode growth media (NGM) plates seeded with *Escherichia coli* OP50.

#### Generation of barcoded lineages

To create the background mutant strain with the TARDIS barcode landing pad, JD608 *avr- 14*(ad1302)I; *avr-15*(ad1051)*glc-1*(pk54)V was crossed with PX740 N2-PD1073 fxIs47[*rsp-0p*:: 5’ Δ*HygR*:: GCGAAGTGACGGTAGACCGT :: 3’ Δ*HygR*::*unc-54* 3’::LoxP, II:8420157] II:8420157 to create PX776 *avr-14*(ad1302)I; fxIs47; *avr-15*(ad1051) *glc-1*(pk54)V. PX776 was injected with TARDIS barcodes following the protocols of Stevenson et al., 2023, with a unique barcode sequence ‘NANNNTNTNNCNNNN’ to facility correct identification of the mutant by sequencing, resulting in PX787 *avr-14*(ad1302)I; fxIs47; *avr-15*(ad1051) *glc-1*(pk54)V; fxEx29 [TARDIS 5′Δ*HygR*::Intron5’::Read1::NANNNTNTNNCNNNN::Read2::Intron3’::3’ Δ*HygR hsp- 16.41p*::*piOptCas9*::*tbb- 2* 3 ’UTR+rsp-27p::*NeoR*::*unc-54* 3’ UTR+U6p:: GCGAAGTGACGGTAGACCGT]; fxIs47. For the wildtype barcoded TARDIS array, PX786 was used and described in Stevenson et al., 2023. Lineages were generated from both PX786 and PX787 following standard TARDIS-integrated protocols (Stevenson et al., 2023). Briefly, TARDIS-array bearing strains were hypochlorite synchronized, and heat shocked at the L1 stage to integrate barcodes, marking the lineage. Several lineages were isolated and identified by Sanger sequencing (Azenta Life Sciences, South Plainfield, NJ).

#### Experimental evolution, liquid culture, and sample collection

To create our liquid environment, we used NGM buffer as our base (Leung et al., 2011), in addition, we added 100µg/ml carbenicillin, 5µg/ml cholesterol, 125µg/ml hygromycin B, and 10µg/ml nystatin. A 10µM ivermectin/DMSO stock was diluted further with DMSO to achieve the desired experimental molarity. DMSO only was used for all the controls and all [DMSO] (including the controls) were normalized for each experimental set while maintaining a final [DMSO] of ≤1% (AlOkda and Raamsdonk, 2022). 4×10^9^ PXKR1 cells/ml (NA22 transformed with pUC19 for carbenicillin resistance) were also added (Stevenson et al., 2023). Bacteria were grown in several large batches and measured for cell concentration before being frozen at -80°C until needed. Independent lineage populations were started by allowing large density plates (100mm) to reach starvation and then each lineage was added independently into a liquid solution. Lineages were then mixed for the parental generation at approximately 10% wildtype, 90% mutant for ivermectin concentrations of 0nM; and 30% wildtype, 70% mutant for1nM; while for 2nM, 3nM, 4nM, and 5nM, populations were mixed at approximately equal concentrations. Each parental population was started with several thousand individuals (supplementary data file 1). Serial cultures were grown in 300ml volumes in 2L flasks mixed with magnetic stir bars and 10% of the population by volume was transferred every five days.

Cultures were maintained at a constant 20°C in a temperature-controlled room (supplemental figure 1). Population densities were estimated by counting six individual drops ranging from 2- 20µl on a glass slide. In some cases, a 10X dilution was made to simplify the counts. Several 1ml samples were taken on the day of transfer and frozen at -20°C. In cases of lower population densities, 10-50ml samples were taken and centrifuged to create a pellet to ensure extra genomic DNA could be acquired. Samples were then processed for genomic DNA and barcode frequency as described in Stevenson et al., 2023.

#### Fitness Estimations and analysis of data with FitSeq

Barcode frequencies derived from Illumina sequencing (see Stevenson et al., 2023) were provided to PyFitSeq–a python implementation of FitSeq (Li et al., 2018). Briefly, FitSeq requires the user to provide the approximate generation time per transfer, which was approximately one generation per transfer, along with estimated population sizes. Only mutant lineages which survived to the end of the experiment–fitness greater than -1– were counted.

Mutant lineages counts with ten or less were excluded from the analysis. Mutant selection coefficients were normalized to the average wildtype fitness. Barcodes which did not confirm to the following sequence ‘NANNNTNTNNCNNNN’ for mutant and ‘NNNCNNTNTNANNN” for wildtype were excluded.

#### Probability of reaching adulthood during ivermectin exposure

Individual barcoded lineages PX905 (wildtype background) fxSi67 [*rsp-0p*:: 3’ Δ *HygR*::Intron5’::Read1::GAGCAATTTAATCAT::Read2::Intron3’::*unc-54* 3’::LoxP, II:8420157] and PX925 (mutant background) avr-14(ad1302)I; fxSi66 [*rsp-0p*:: 3’ Δ *HygR*::Intron5’::Read1::GATAATATACCTAGT::Read2::Intron3’::*unc-54* 3’::LoxP, II:8420157]; *avr-15*(ad1051*) glc-1*(pk54)V were grown on plates until the population contained mostly gravid adults. The populations were then synchronized in NGM buffer by bleaching the adults in a solution of 1% sodium hypochlorite/0.5% NaOH to collect the eggs. For each strain, eggs were counted post-synchronization (as described above), and liquid culture solutions were made which contained one egg/µl. Cultures were then exposed to DMSO or DMSO with concentrations of ivermectin in a liquid culture solution as described above, except they were grown in 15ml conical tubes and allowed to rotate at 20°C to ensure proper mixing and aeration. Total liquid volumes were 5ml for each culture to allow substantial air space within the tubes.

Just prior to counting, populations were immobilized with 0.2mM levamisole. For each timepoint, several 20µl drops were scored for both the total number of animals and the number of animals that had reached the adult stage at two separate timepoints, 72 hours and 96 hours post synchronization, to obtain the percentage adults. Animals were determined to be adults if gravid (eggs observed) or if no eggs, by examining individual animals for mature vulva development at the highest magnification (112.5X).

#### Microscopy

600µl samples from the developmental liquid cultures were centrifuged and 550µl of the supernatant was removed to create a denser population for imaging (∼50 µl). 3µl of each worm concentrate was then placed onto a glass slide and cover slipped. Imaging was performed on an Olympus IX73 using cellSens software v2.3. Samples were imaged under white light for 20 milliseconds exposures using a 4x objective. Scale bars were added using Fiji (imageJ) v2.9.0/1.53t.

#### Accessibility of reagents, data, code, and protocols

The authors affirm that all data necessary for confirming the conclusions of the article are present within the article, figures and tables. Plasmids pZCS36 (Addgene ID 193048), pZCS41 (Addgene ID 193050) are available through addgene and can be freely viewed with ApE (Davis and Jorgensen, 2022). Strains are available upon request. Illumina sequencing data is available at NCBI BioProject ID: PRJNA1170954. All barcoding count information, adult counts, and fitness data are available in supplementary file one. Original images of ivermectin exposed worm for qualitative developmental assessment is available in supplementary file two. All statistical analysis and code are available in supplementary files three and four.

#### Software and statistical analysis

Lineage frequencies were visualized with matplotlib 3.5.2 and data was analyzed with Python 3.7.13. For selection coefficient, peak census size, and developmental trade-offs, plots and statistics were generated in R v.4.3.2 (Team, 2023), lmer version v.1.1.35.3 (Bates et al., 2015), and visualized using ggplot2 v.3.5.1 (Wickham, 2016). Least means squared was calculated using the emmeans package v. 1.10.1. All code was executed in either Jupyter Notebooks v3.6.3 (Google Colab)–stacked frequency plots, or Jupyter Labs v7.9.0 (Kluyver et al., 2016) –all statistics done in R, along with plots for the selection coefficients and developmental trajectories.

## Acknowledgments

We thank the Phillips lab members for helpful suggestions and technical assistance. In particular, we thank Megan J. Moerdyk-Schauwecker and Stephen A. Banse for their assistance and general discussion on the molecular biology, Christine Sedore for her assistance in statistical analysis. We would also like to thank Runpeng Nie and Julia Hibbard for their assistance in the experimental evolution. Some strains were provided by the CGC, which is funded by NIH Office of Research Infrastructure Programs (P40 OD010440). We also want to acknowledge and thank WormBase for providing a database resource of strains and genes used in this study.

## Description of supplementary files

Supplementary Data File 1-Count Data For Fitness Development Census and Index Associated for Demultiplexing

Supplementary Data File 2-Representative Images of Worms Developing in Ivermectin After 72 Hours

Supplementary Data File 3-Fitness Analysis Supplementary Data File 4-Developmental Analysis

